# Improved accuracy for estrous cycle staging using supervised object detection

**DOI:** 10.1101/2024.05.08.593231

**Authors:** Benjamin Babaev, Saachi Goyal, Rachel A Ross

## Abstract

The estrous cycle regulates reproductive events and hormone changes in female mammals and is analogous to the menstrual cycle in humans. Monitoring this cycle is necessary as it serves as a biomarker for overall health and is crucial for interpreting study results. The estrous cycle comprises four stages influenced by fluctuating levels of hormones, mainly estradiol and progesterone. Tracking the cycle traditionally relies on vaginal cytology, which categorizes stages based on three epithelial cell concentrations. However, this method has limitations, including time-consuming training and variable accuracy among researchers. To address these challenges, this study assessed the feasibility and reliability of two machine learning methods. An object detection-based machine learning model, Object Detection Estrous Staging (ODES), was employed to identify cell types throughout the estrous cycle in mice. A dataset of 555 vaginal cytology images with four different stains was annotated, with 335 images for training, 45 for validation, and 175 for testing. A novel, accurate set of rules for classification was derived by analyzing training images. ODES achieved an average accuracy of 87% in classifying cycle stages and took only 3.9 minutes to analyze 175 test images. The use of object detection machine learning significantly improved accuracy and efficiency compared to previously derived supervised image classification models (33-45% accuracy) and human accuracy (66% accuracy), refining research practices for female studies. These findings facilitate the integration of the estrous cycle into research, enhancing the quality of scientific results by allowing for efficient and accurate identification of the cycle stage.

## INTRODUCTION

### The Estrous Cycle

The estrous cycle, analogous to the human menstrual cycle, is the reproductive cycle of female non-primate vertebrates including mice and rats. The estrous cycle is divided into 4 stages: diestrus, proestrus, estrus, and metestrus. These stages are influenced by fluctuating levels of ovarian hormones, notably estradiol and progesterone. The cycle influences brain activity [1] and physiological characteristics, such as changes in hypothalamic-pituitary-adrenal axis activity [2], feeding, and energy metabolism [3]. However, the effects of the cycle are often overlooked in neuroscience investigations and psychiatric care [1]. Historically, research has predominantly utilized male subjects, resulting in an insufficient understanding of female disease physiology and suboptimal treatment plans for females [4]. The common misconception that data/results from female studies are more variable than groups of males has not been proven true. Instead, it seems that there are fundamental differences as a result of sex-based variation [5], including some that are due to hormone fluctuations across the cycle. Characterizing the stages of the estrous cycle serves as a tool for understanding the female biological context [6]. Incorporating the estrous cycle in studies related to rodent behavior and genetic interactions can improve the quality of neuroscience findings and more accurately identify hormone-fluctuation-related sex differences. Recognizing hormone variance is crucial in examining behavior, pharmacology, and neurological changes, and drawing parallels between animals and humans provides an opportunity to understand the mechanisms that underlie these changes.

### Estrous Stage Classification Techniques

The identification of the estrous stage is typically done using vaginal cytology, visual assessment of the vagina, or vaginal wall impedance [7]. Vaginal cytology is the most common indirect tracking method and involves microscopic examination of vaginal smears, analyzing the shape, size, and number of cells to determine each stage. Epithelial cells are found in the vaginal lining and change in appearance throughout the estrous cycle. Cornified cells, which lack a clear nucleus, are most abundant during estrus which lasts 12–48 hours. Nucleated epithelial cells are present in higher numbers during proestrus which lasts less than 24 hours. Leukocytes, or white blood cells, are most abundant during diestrus which lasts 48–72 hours. The metestrus stage shows a mixed cell population with no distinct majority (Figure 1) and lasts 8–24 hours. Because the hormone ratios are similar and it can be difficult to differentiate by cytology, metestrus is also commonly combined with diestrus, or referred to as early diestrus.

**Figure 1.**
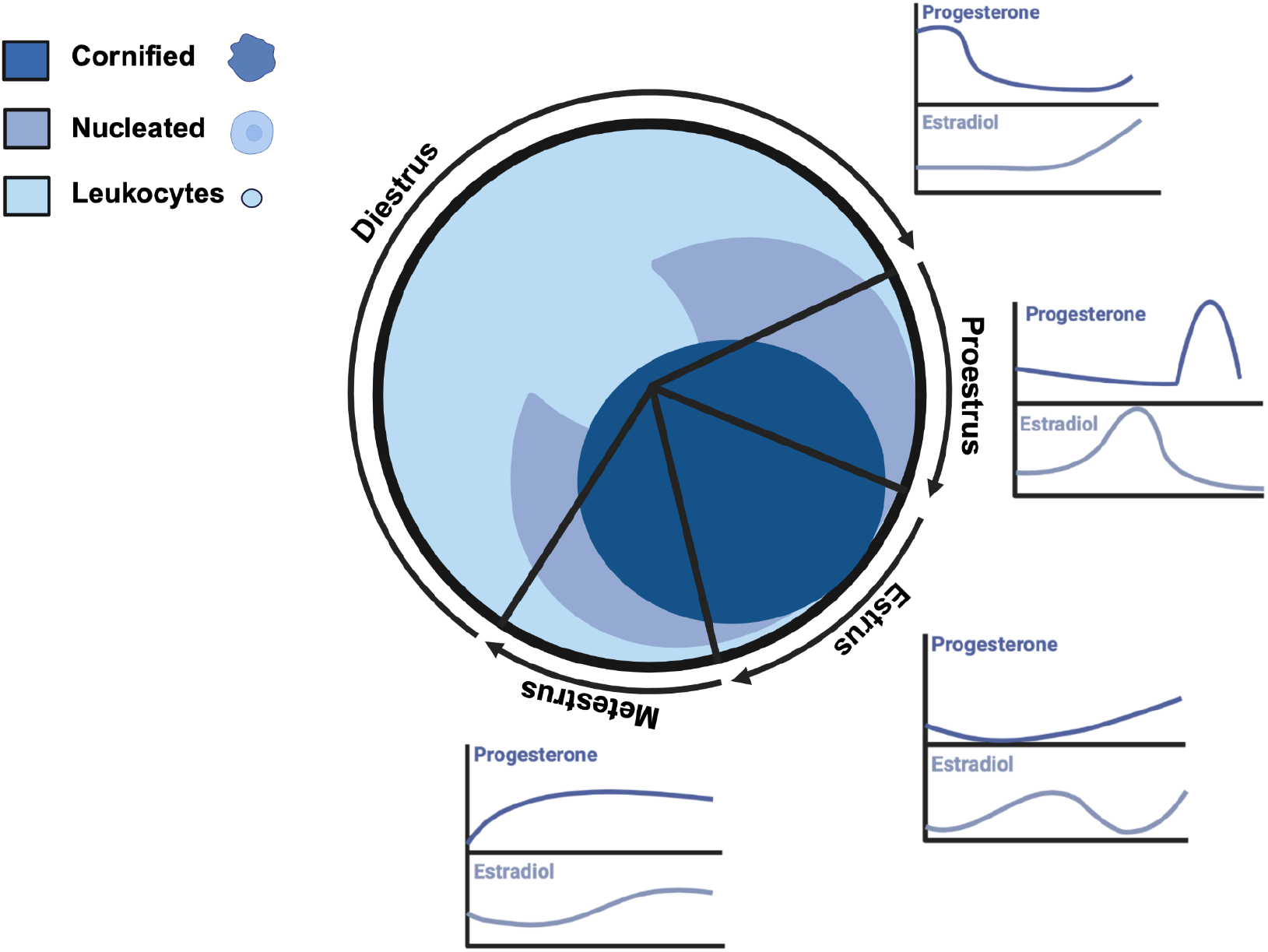
Cell populations and relative sex hormone levels throughout the estrous cycle in female mice (Adapted from Byers et al., 2012 and Donner & Lowry, 2013). Schematic made with BioRender.

### Drawbacks of Vaginal Cytology and Previously Used Machine Learning Strategies

Vaginal cytology has limitations: it requires significant time and training to ensure proper classification, and there can be variability in classification among researchers. To overcome these drawbacks, previous research has explored the application of image classification. These supervised machine-learning techniques utilize convolutional neural networks which have proven to be more accurate and efficient than human experts at diagnosing various medical disorders like cancer, heart disease, and neurological conditions [8]. The ability to do so arises from analyzing vast amounts of data, which allows for the identification of patterns and anomalies that may be overlooked by human specialists. However, a limitation of image classification is the inability to identify the location of various objects within an image, therefore the exact rationale behind the model’s stage classifications is unknown [9].

Two supervised image classification models for vaginal smears have been previously described. The SECREIT model [10] was developed to identify images representing three stages of the estrous cycle—diestrus, estrus, and proestrus—using solely Giemsa stain, achieving a reported accuracy of 93%. The EstrousNet Model [11] was trained on thousands of images from various labs, encompassing different animal species (mice and rats), stains, and magnifications, with a reported accuracy of 83%. However, it is yet unknown whether the accuracies of these models are reproducible on new data sets.

### Object Detection

Supervised learning is a common form of machine learning that can create models to accurately predict outcomes for new, previously unseen data when trained on large labeled datasets [12]. It is widely used for tasks such as image classification and object detection. Image classification involves assigning a single label to an entire image. Object detection, on the other hand, involves the identification and localization of objects in a given image. This technique is useful for its ability to detect objects across different classes, sizes, orientations, and scales [13]. Its advantages over image classification are that it recognizes both the presence and position of various objects, each with a respective label and that it has the capacity to combine information from an array of verified objects in order to classify the image. It requires human input to define individual objects, which can be a source of bias, but this is mitigated in the context of a limited object dataset such as cells seen in vaginal smears.

YOLOv8, short for “You Only Look Once” version 8, is a freely available object detection model that uses the principles of convolutional neural networks to perform real-time object detection with high precision. We used the YOLOv8 model to achieve generalizability and accuracy in classifying vaginal cytology images, surpassing both image classification models and human experts. This standardized, rule-based approach minimizes variability among researchers and reduces classification time. Here we demonstrate the potential of a supervised object detection paradigm called Object Detection Estrous Staging (ODES) in automating cell type classification and cell counting, offering a method similar to human expert classification of vaginal cytology, with enhanced efficiency and generalizability.

## METHODS

### Dataset and Preparation

The dataset for this study was composed of mouse vaginal cytology images sourced from the Ross Lab, Verstegen Lab, Correa Lab, Sano Lab, and EstrousBank, a free online resource with images from five different labs [11]. The dataset staining techniques included the Giemsa stain, Crystal Violet, Cresyl Violet, Shorr stain, and H&E stain, and multiple magnification levels. The data was divided into three sets of randomly chosen images: 335 images for training, 45 for validation, and 175 for testing.

### Image Annotation

To prepare the dataset for training, the images were annotated using a freely available online labeling tool makesense.ai. Each image in the training set was manually annotated by drawing rectangular bounding boxes around each cell and tagging the cell with its corresponding label: “Leukocyte”, “Cornified”, or “Nucleated”. The training set had 9226 instances of leukocytes, 12584 instances of cornified cells, and 4349 instances of nucleated cells (Figure 2a).

**Figure 2.**
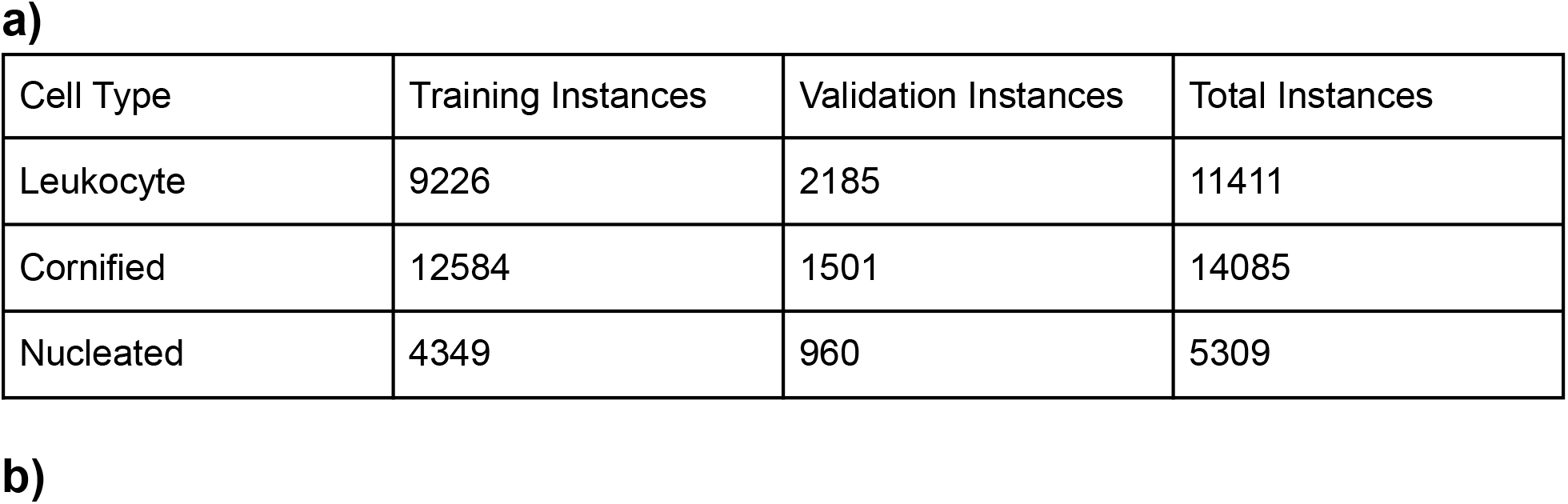
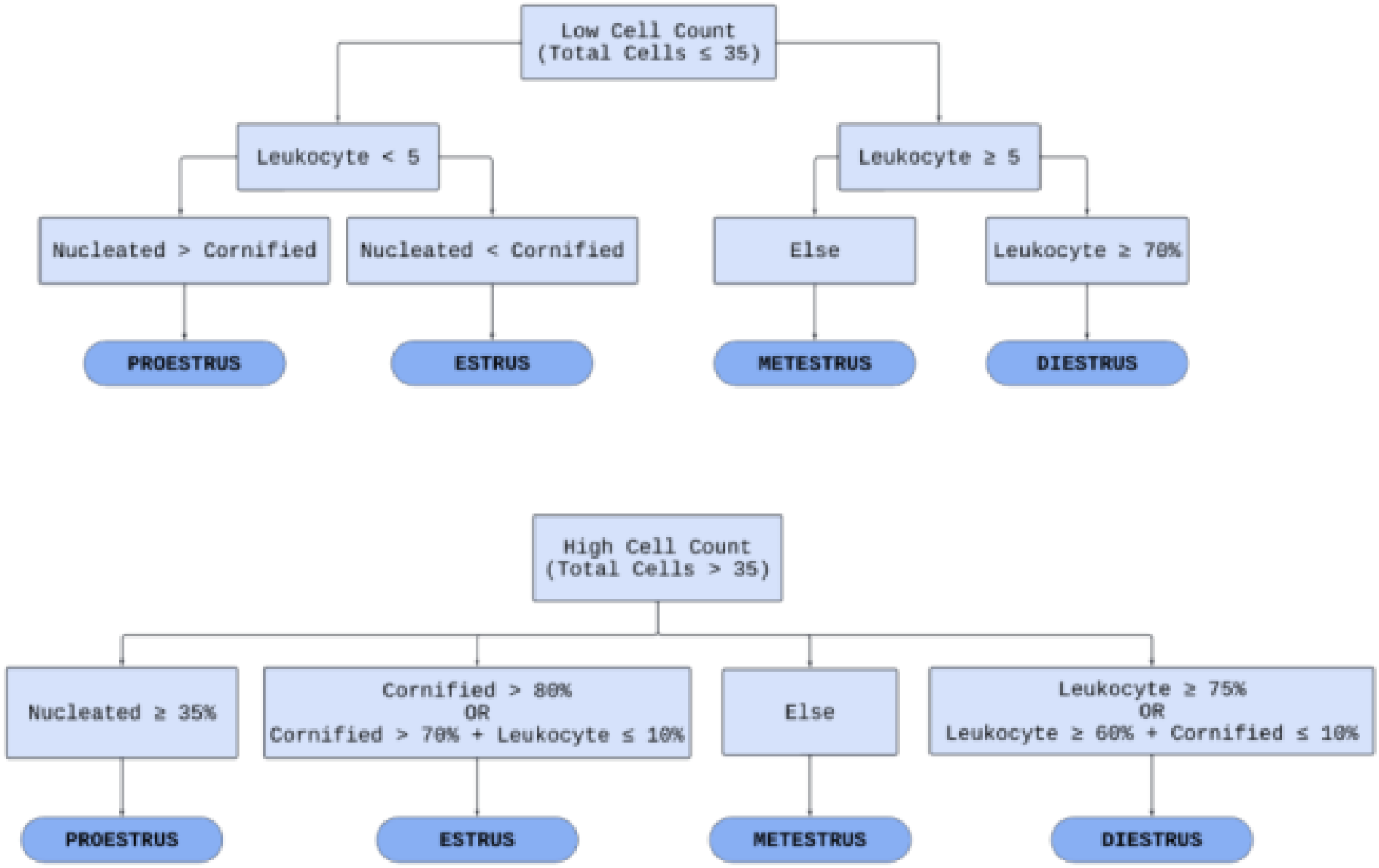
**a:** Table displaying the distribution of cell types across the training and validation datasets. The leukocyte class consisted of 9,226 instances in the training set and 2,185 in the validation set, with a total of 11,411 instances. The cornified class consisted of 12,584 instances in the training dataset and 1,501 in the validation dataset, with a total of 14,085 instances. The nucleated class had 4,349 instances in the training set and 960 instances in the validation set, resulting in a total of 5,309 instances. **b:** Flowchart that is a visual representation of the algorithm (written in Python) used to determine the estrous stage by assessing the cell count. It divides the process into two algorithms: one for low cell count and another for high cell count. Through a series of binary decisions based on the criteria provided in each block, the model narrows down the classification to one of the four estrous stages.

### Exporting Annotated Images

Post-labeling, the annotations of the images were exported as .txt files. We used a class index for each cell type: 0 for Leukocyte, 1 for Cornified, and 2 for Nucleated. Additionally, the annotation included four unique values to represent the x-center coordinate, y-center coordinate, and width and height of the bounding box [14]. The raw images were downloaded into the image folder and the corresponding YOLO text document, with the same name as their corresponding image, into the labels folder. The images with their corresponding .txt files were sorted randomly into training and validation folders.

### YOLOv8 Architecture and Features

The backbone of the YOLOv8 object detection framework is an advanced version of the CSPDarknet53 backbone, which reduces processing time and memory usage [15,16]. CSPDarknet53 is an advanced neural network architecture that focuses on efficient feature extraction from input images [17]. It employs the Cross Stage Partial (CSP) design to balance detail and efficiency, improving the network’s ability to capture important visual details and broader scene context [18,19]. The model includes a Spatial Pyramid Pooling (SPP) Fast layer to improve detection accuracy and speed by optimizing feature extraction [16]. SPP divides input feature maps into regions of various sizes, enabling the network to capture details at multiple scales essential for accurate object detection [20]. It processes information sequentially by pooling kernels of the same size rather than analyzing all sizes simultaneously [20]. Other features like spatial attention improve the accuracy of object detection by enhancing the model’s ability to recognize objects in complex backgrounds [16].

### Model Training

The pre-trained YOLOv8s model (a small version of YOLOv8) was utilized within a Google Colaboratory environment, which provided access to the Tesla T4 GPU. The model was adapted to recognize specific classes of cells (leukocytes, cornified cells, nucleated cells) not included in its original training set. This was achieved by employing a custom dataset configuration to guide the model’s training process. The document “customdata.yaml” provided explicit paths to the training and validation sets, and introduced the three cell-type classes with their respective class indices. Images of various sizes were used for training and validation but for YOLOv8, the image-size parameter was set to 640. The model was trained for 62 epochs, during which various augmentations were applied to diversify and enlarge the training set. The default augmentations utilized by YOLOv8 were applied, including changes in hue, saturation, brightness, translation, scaling, horizontal flipping, mosaic augmentation, random color and rotation variations, and random erasing. The maximum object detection parameter was set to 1000. The batch size was set to 18 and the patience was set to 20 epochs to prevent overfitting. The weights of the trained model were then saved to allow for testing.

### Rule-Based Classification

A preliminary rule-based stage classification system (Figure 2), written in Python, was developed using a qualitative diagram of the estrous cycle [21]. The performance of the trained model was evaluated by observing its outputs when presented with novel test images it had not previously encountered. The rules and training criteria were refined through a feedback loop to improve the model’s accuracy and better represent the cell populations of each stage. The feedback loop involved training the model on the training set, evaluating the model’s performance on the validation set, and adjusting either the rules or training criteria based on validation results (Figure 2a). If the model fell short in cell classification, the validation image was annotated, placed into the training set, and retrained. If the model misclassified the estrous stage and the cell classification was accurate, the rule-based stage classification system was adjusted (Supplemental Figure 1). An additional set of rules was developed to account for cases with low cell counts (Figure 2b). For instances where ODES output showed a lower cell count than expected or a higher background count a “Biological impossibility tag” was implemented.

This tag flags scenarios where the cell counts are low, which could indicate that the model failed to detect clusters of cells and instead misclassified them as background. The tag is indicated by three asterisks (***) in the output classification table the user receives upon running ODES on a set of vaginal cytology images. Once the rules and weights were finalized, the model was complete.

### Model Comparison

The performance of ODES was tested against human accuracy. The study protocol was reviewed by the Albert Einstein College of Medicine Institutional Review Board. Four human researchers were timed while classifying a set of 100 images. The model ran through the same set of 100 images and was timed. The test set contained random images obtained from EstrousBank and SECREIT data that ODES had not seen before.

The model was also compared with two previously published supervised image classification machine learning models: EstrousNet and SECREIT. Two separate test sets were utilized: one consisting of 100 images sourced from EstrousBank data, with 25 images representing each of the four estrous stages, and another consisting of 75 images from SECREIT data, with the same stage distribution but excluding the metestrus stage, as their data combined diestrus and metestrus.

The accuracy of the participants and the model was determined by comparing their classifications to the established benchmarks provided by EstrousBank and SECREIT datasets, which were considered the correct stages for this experiment.

### Statistics

A nested one-way ANOVA followed by Tukey’s multiple comparisons test was used to compare the classification accuracy of ODES with EstrousNet and SECREIT across estrous cycle stages. Cohen’s d was used to compare the effect size of the accuracy difference between ODES, EstrousNet, and SECREIT. A two-way ANOVA followed by Sidák’s multiple comparisons test was used to compare classification accuracy and classification time per image between ODES and human experts across estrous cycle stages.

## RESULTS

### ODES Cell Classification

The trained model’s cell classification ability was analyzed on the 45-image validation set. The performance is demonstrated with a confusion matrix, providing the prediction accuracy for each cell type: leukocyte, cornified, and nucleated (Figure 3a). Leukocytes were correctly identified in 93% of cases with only a 6% misclassification rate with nucleated cells and 1% as cornified cells. Cornified cell identification was 90% accurate, with 10% of instances being mistaken for nucleated cells. The accuracy for nucleated cells was 72%, with misclassification as cornified cells to 15% and leukocytes at 13%. In this classification, many cells, particularly those that overlap and clump together, were mistaken for background (Supplemental Figure 2). This was primarily the case for leukocytes which tend to be clustered, posing challenges for ODES in distinguishing them from background elements. To mitigate this issue, a ‘Biological Impossibility Tag’ was introduced to flag low cell counts that may suggest the presence of uncounted cell clusters or other anomalies. It is recommended that users using ODES review the images with low cell counts.

**Figure 3.**
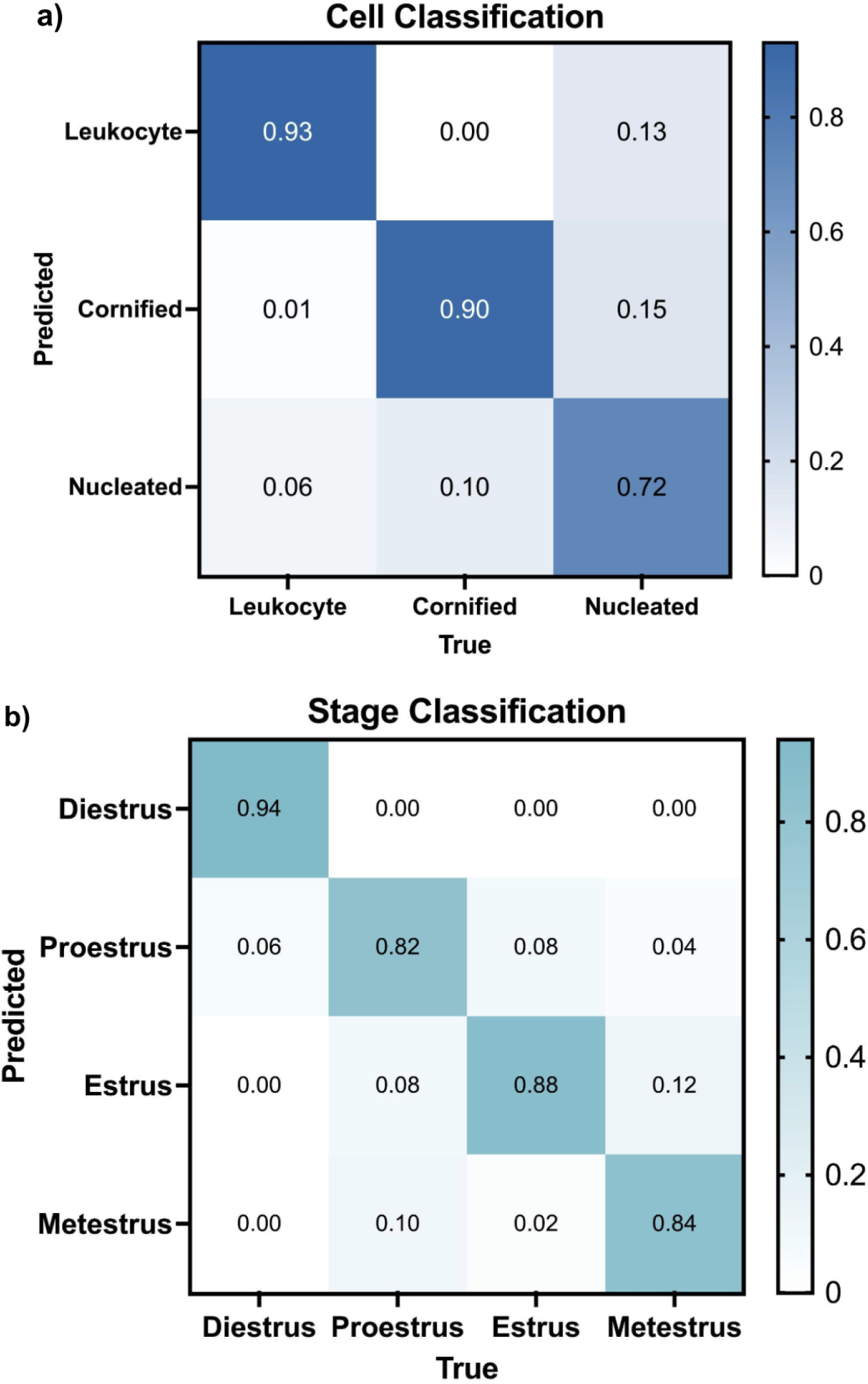
**a:** Normalized confusion matrix that evaluates the cell classification performance of ODES on the validation set. The x-axis represents the true cell type (Leukocyte, Cornified, Nucleated), and the y-axis shows the predicted labels. The diagonal elements of the matrix represent the instances that were correctly classified by the model, while the other elements represent the number of instances that were misclassified. **b:** Normalized confusion matrix for ODES stage classification of 175 test images (50 diestrus, 50 proestrus, 50 estrus, 25 metestrus). The x-axis indicates the true stages (Diestrus, Proestrus, Estrus, Metestrus), while the y-axis shows the predicted stages. Similar to Figure 3a, the diagonal represents the correct predictions for each stage and the other cells represent the incorrect predictions.

### ODES Stage Classification

Using the classification rules (Figure 2b) and trained weights derived from cell classification training, ODES classified the stages of the estrous cycle in a test set comprising 175 previously unseen images (100 from EstrousBank, 75 from SECREIT). The stage classification confusion matrix demonstrates the accuracy of the model for each estrous stage (Figure 3b). In the diestrus stage, ODES achieved an accuracy of 94% with misclassification as proestrus in 6% of instances. The true proestrus stage was correctly predicted with an accuracy of 82%, though 10% of instances of proestrus were misclassified as metestrus and 8% as estrus. The estrus stage had an accuracy of 88%, with confusion between estrus and proestrus in 8% of instances. The metestrus stage had an 84% correct classification with misclassification as estrus in 12% of instances and proestrus in the remaining 4%.

To test if any stain provided greater accuracy than others, we evaluated outcomes for each stain type separately. The test set of 175 images was used to test stain accuracy out of which 89 had the Giemsa stain (14 from EstrousBank and 75 from SECREIT), 29 had the H&E stain, and 57 had the Shorr stain. ODES achieved an accuracy of 92.1% for Giemsa, 82.6% for H&E, and 80.7% for Shorr stain. The trend toward best performance on images with a Giemsa stain was not statistically significant (Supplemental Figure 3).

### Comparison of ODES with Image Classification Models

The stage classification accuracy of ODES was compared to two supervised image classification machine learning models, EstrousNet and SECREIT, across the four stages of the estrous cycle (Figure 4a). The test set contained randomly chosen images for each estrous stage obtained from EstrousBank (100 images, 4 stages) and SECREIT data (75 images, 3 stages). For the diestrus stage, ODES achieved an average classification accuracy of 94%. In contrast, EstrousNet and SECREIT had significantly lower accuracies of 74% (p<0.021) and 56% (p<0.028), respectively. In the proestrus stage, ODES’s accuracy averaged 82%, while EstrousNet classified with 18% accuracy, and SECREIT with 10%. During the estrus stage, ODES had an accuracy of 88%, with EstrousNet at 36% and SECREIT at 42%. For the metestrus stage, ODES had an average accuracy of 84%, and EstrousNet’s accuracy was 16%. SECREIT’s performance in the metestrus stage was not assessed as it was not trained to classify metestrus. Taken together, ODES shows significantly improved accuracy and generalizability over the image classification models for vaginal cytology stage classification.

**Figure 4.**
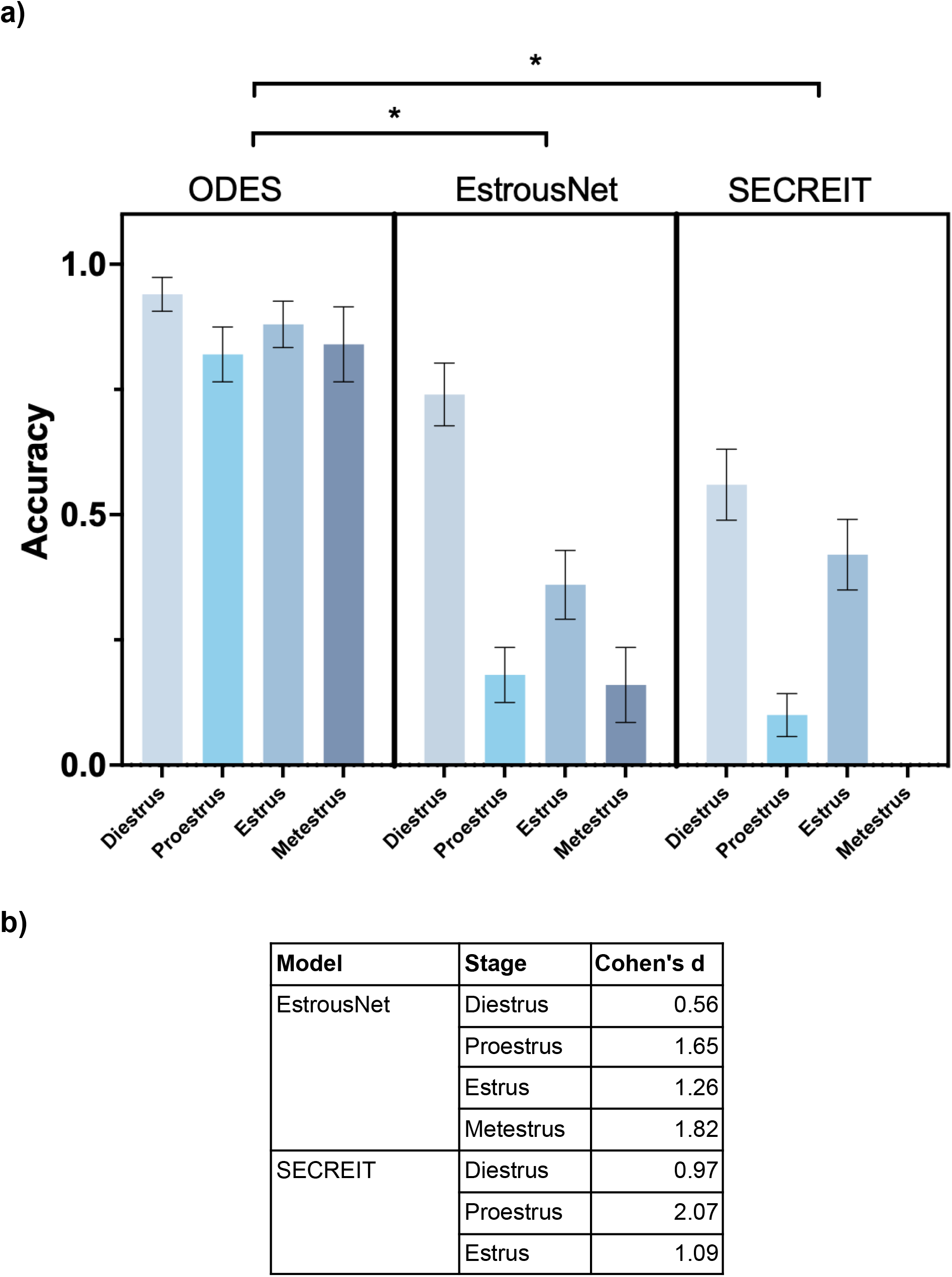
**a:** Classification accuracy of ODES compared to image classification ML models EstrousNet and SECREIT across the four estrous cycle stages. SECREIT was not trained to classify the metestrus stage, therefore data for SECREIT metestrus is not available. A nested one-way ANOVA revealed significant differences in model performance across the estrous stages (p<0.014). Tukey’s multiple comparisons test indicated significant differences between the performance of ODES and EstrousNet (p < 0.021) and ODES and SECREIT (p<0.028). **b:** Table comparing the effect sizes of ODES to EstrousNet and SECREIT on the accuracy of estrous stage classification using Cohen’s d.

In assessing the performance of the ODES model compared to EstrousNet and SECREIT across various estrous phases, Cohen’s d values offer a quantitative measure of the effect size. This effect size analysis reveals that ODES outperforms the other two models across all stages (Figure 4b). For the diestrus phase, ODES shows a moderate improvement in accuracy over EstrousNet with a Cohen’s d value of 0.56, and a large improvement over SECREIT, with a d value of 0.97. In the proestrus phase, ODES shows a large effect size compared to EstrousNet with a Cohen’s d of 1.65. Against SECREIT, the effect size is larger at Cohen’s d of 2.07. In the estrus phase, ODES performs better with Cohen’s d of 1.26 compared to EstrousNet and a similarly large effect of 1.09 against SECREIT. In the metestrus phase, ODES has an effect size of 1.82 against EstrousNet. This quantitative analysis suggests that ODES is a more reliable and effective model for determining the correct estrous phase in images.

### Comparison of ODES to Human Experts

The accuracy rates for both ODES and human classifiers were compared across the four estrus cycle stages (Figure 5a). ODES achieved high accuracy rates with averages of 96% for diestrus, 80% for proestrus, 88% for estrus, and 84% for metestrus. In comparison, human accuracy rates were measurably but not significantly lower, with an average of 56% for both diestrus and proestrus, 79% for estrus, and 75% for metestrus. The average classification time per image was also compared between human experts and ODES (Figure 5b). The human experts averaged 9.7 seconds per image, while ODES reduced the classification time to an average of 1.6 seconds per image (p<0.001). Thus, ODES preserves accuracy and enhances efficiency of vaginal cytology based estrous cycle staging compared to human experts.

**Figure 5.**
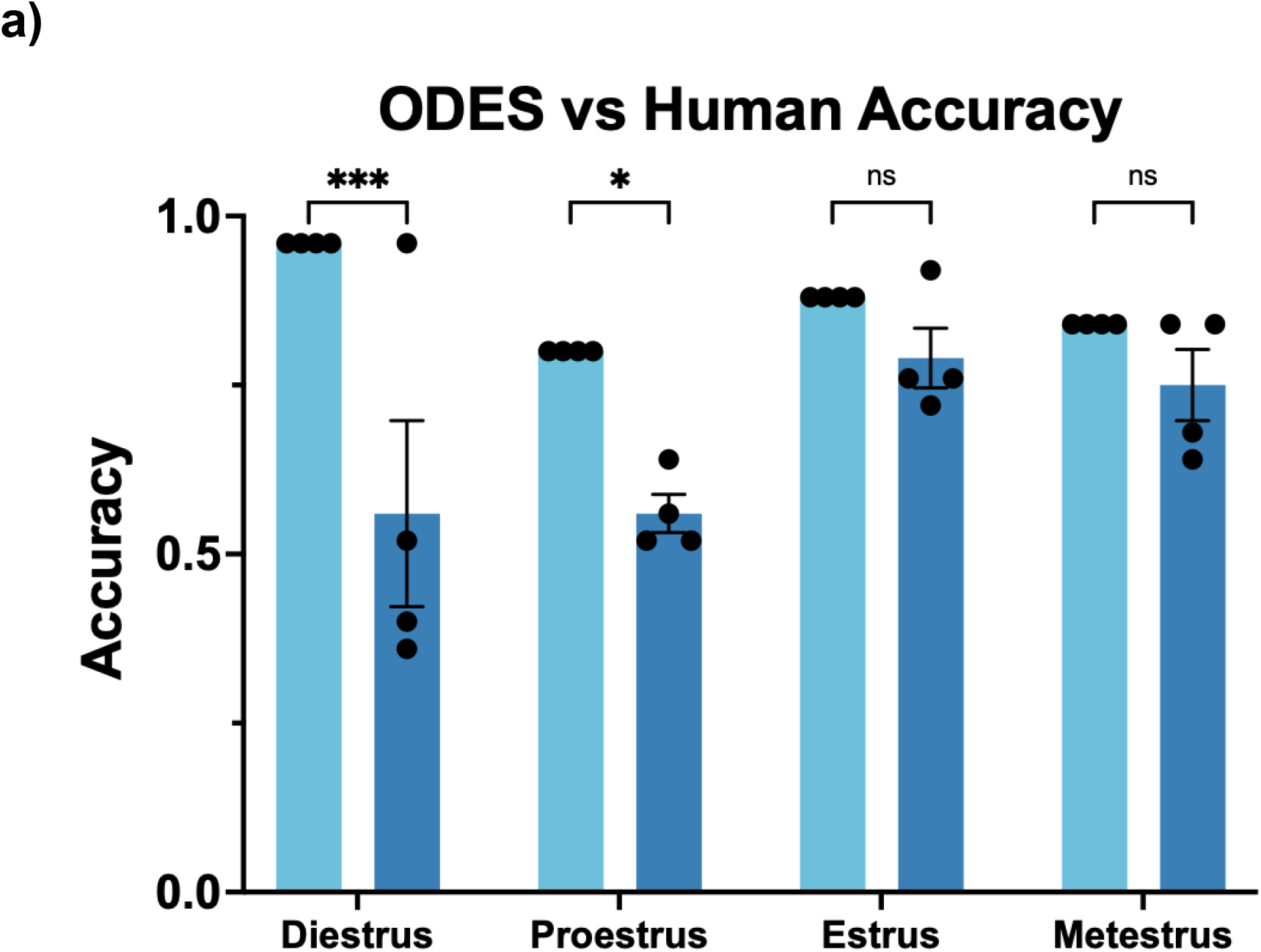
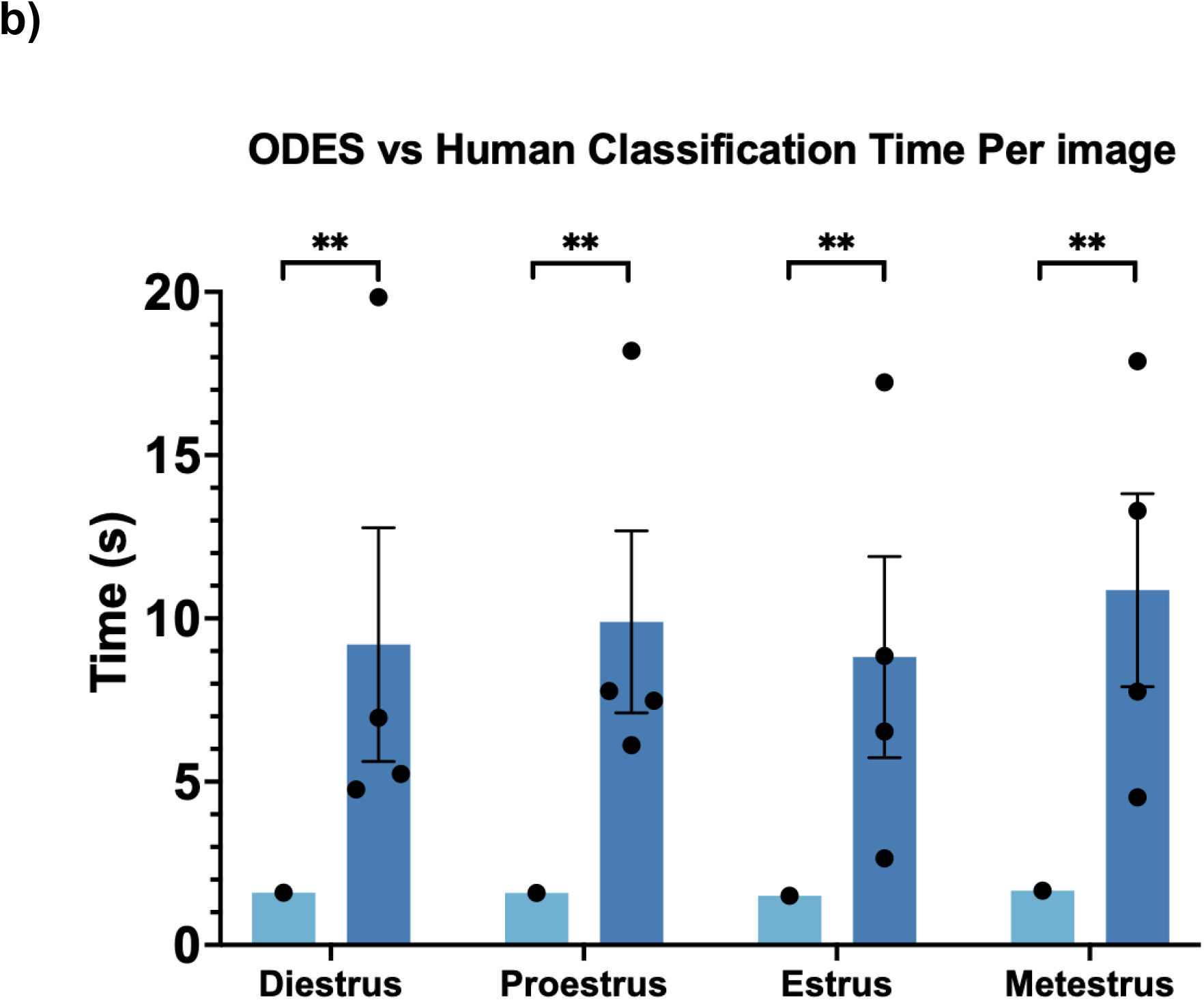
**a:** Comparison of ODES and human accuracy in estrous cycle stages classification. The error bars indicate the SEM, while individual data points represent different trials. ODES was run four times to statistically compare to four human trials. In a two-way ANOVA, there was a significant difference between ODES and Human accuracy (p < 0.0001). Using Sidák’s multiple comparisons test, significant differences were present for diestrus and proestrus, with p-values of 0.0010 and 0.0384 respectively, however there were no significant differences in estrus and metestrus, both with p-values of 0.7195. **b:** Comparison of ODES and human classification time per image in estrous cycle stages classification. The error bars indicate the SEM. The individual data points represent the time per image averaged over 25 images. In a two-way ANOVA, there was a significant difference between ODES and Human time (p < 0.001). Using Sidák’s multiple comparisons test, significant differences were present for all four stages (p<0.0015).

## DISCUSSION

We used object detection to train a machine learning model to approach images of vaginal cytology like an expert scientist. We trained the model to recognize individual cell types and created a rule scheme that allowed the model to define the estrous cycle stage based on the relative percentage of each cell type seen in each image. ODES demonstrates its classification capabilities by outperforming image classification models in generalizability and accuracy, and human experts in classification speed. It has also shown consistent accuracy across various stains, magnifications, and data sources. The higher performance of ODES in classifying estrous stages may be attributed to its novel approach that integrates object detection with subsequent classification based on detected cell types. This methodology provides a contrast to traditional image classification models that rely only on supervised learning paradigms [10,11]. Previous models typically label entire images according to the estrous stage, without the ability to identify individual cell types [10,11]. This may have led to pattern detections within the image that do not capture the features essential for accurate stage classification.

By identifying specific cell types and quantifying their proportions, ODES’s approach allows for a more detailed and biologically relevant analysis. There is a well-defined, limited data set that includes all possible cell types that would be present in a given vaginal cytology slide. Therefore, it is reasonably simple to train the paradigm to recognize all objects in this limited set, and there is minimal bias imposed by human input. The model’s ability to match cellular features to their specific types and then classify them based on these more detailed metrics may have ensured a higher degree of accuracy in stage classification, even with confusion between some cell types and the background. This two-step method allows the model to be generalized and scalable in ways that less supervised, or less detail-oriented models may not.

While the goal of machine learning is generally to reduce human input, it seems that in this case, there is a significant benefit to human involvement in the supervised learning process. A loss function measures the model’s performance by calculating the difference between the model’s predictions and the actual labels [22,23]. The goal is for this “loss” to be as small as possible by adjusting the model’s parameters to minimize this difference between predicted and actual labels. The loss function which can be applied to supervised learning ultimately improves the accuracy of the model’s predictions. Additionally, the supervised model is designed to minimize generalization loss, ensuring reliable performance on unseen data (such as the test set). This is important in cell classification where sample variability can be high but remains advantageous because the set of objects is limited by biology. Future work should evaluate whether ODES accuracy is maintained in a new context, such as a different stain type, or a dataset from another animal model.

### Challenges with specific cell types and stages

All models, including ODES, had the lowest accuracy of stage classification for the proestrus stage. Similarly, ODES’s cell classification accuracy for nucleated cells, which are the predominant cell type during proestrus, was the least accurate. There are a couple of reasons for this poor performance. First, the limited duration of the proestrus stage, typically less than 24 hours, led to a smaller dataset for this phase. Smaller data sets often lead to imbalanced data and model overfitting or underfitting due to inappropriate feature dimensions, a common challenge in machine learning [24]. The limited data availability therefore restricts ODES’s ability to accurately classify nucleated cells, affecting the overall classification performance for the proestrus stage. To address this, a redefined threshold for proestrus classification is suggested in the stage classification flow chart: a nucleated cell percentage of 35% or higher, instead of a simple majority as initially encoded based on Byers et al [21]. This increased the accuracy of ODES proestrus readout such that of all models and humans tested, ODES has the highest average accuracy for proestrus staging.

### Limitations of ODES

ODES has not yet been trained to recognize unstained images, which could be seen as a limitation as this adds an extra step for human researchers. Additionally, as previously mentioned, ODES has difficulties classifying individual cells when they are clumped together, and classifying stages when cell count is very low. The difficulties with clumped cells arise due to the lack of clear boundaries between the cells. Cells can overlap each other, creating complex morphological features that are challenging to distinguish. Since ODES was trained to make decisions in a manner similar to human judgment, it encounters the same problems with clumped cells that humans do. Additionally, since the model’s training involves human input, any bias or inconsistency in cell annotations can directly influence the model’s learning patterns [25]. For instance, if the annotations provided by human experts vary due to subjective interpretations of what constitutes a particular cell type or boundary, the model will learn these inconsistencies, potentially reducing its accuracy and generalizability. Additionally, human annotators can have varying levels of confidence and criteria in classification [25]. This variability can lead to a model that is uncertain or inconsistent in its classifications, mirroring the inconsistencies of its training data.

In assessing the performance of ODES, it is also important to consider the data used, particularly the stage labels provided by various labs. There is a possibility that these labels may not have been completely accurate. This potential mislabeling in the source data can affect the accuracy of the model by introducing a source of error that is external to the model’s logic and training process. Future work might involve validating the dataset labels.

Additionally, while it’s common practice to classify estrous images by considering their order [21], ODES currently does not incorporate data in a particular sequence. Integrating this logic, potentially by standardizing the order of the dataset, could improve accuracy outcomes, especially across data collected over time from individual animals.

### Insights from supervised classification at different levels

ODES’s higher accuracy in stage classification compared to cell classification offers valuable insights into stage determination by the model. It seems that the primary requirement of stage classification accuracy is to correctly identify the dominant cell type within the image a sufficient number of times, rather than achieving perfect accuracy for every cell. As a result, misclassifications of individual cells have a reduced impact. If the model can accurately classify a majority of cells correctly, the true stage determination is likely to be achieved. This principle is seen in other studies as well. For example, a study using phase imaging with computational specificity has shown that even with large sample sizes and varying cell counts, accurate cell stage classification is achievable [26]. This reinforces the idea that classifying the majority of cells correctly can be sufficient for accurate results, suggesting that ODES is likely to be similarly scalable.

The cell classification confusion matrix with background (Supplemental Figure 2) shows that most of the error in cell classification occurs when the cells are mistaken for background. Upon review of these images, we found this is due to challenges in recognizing clusters of cells. The difficulty seems to lie in ODES’s ability to recognize and differentiate the boundaries of overlapping individual cells. In these instances, ODES output showed a lower cell count than expected or a higher background count, both of which led to the implementation of the “Biological impossibility tag”, which alerts users that the image may be misclassified. By flagging potential errors, we introduce a secondary review process: both manual check by a human expert and classification through another automated rule set designed to classify images with low cell count. This approach enhances the overall accuracy and reliability of the system.

### Future Directions for ODES

Future improvements for ODES focus on two main areas: expanding the dataset and upgrading the model architecture. The inclusion of unstained images into the training and validation set, and ordered images defined according to individual animals may enhance the model’s accuracy and adaptability to various imaging conditions as well as further improve its generalizability.

The training of ODES was limited in computational resources, such as storage capacity in the training environment, which restricted the ability to employ larger, more complex YOLO models. With access to hardware with greater GPU capacity, it would be feasible to implement larger and more complex versions of the YOLOv8 model. Additionally, with the recent release of YOLOv9, there is potential for improved training results when compared to YOLOv8. This is attributed to its innovative approach to mitigating information loss challenges commonly observed in deep neural networks.

## Conclusions

ODES has demonstrated remarkable generalizability and accuracy in classifying vaginal cytology images, outperforming both image classification models and human experts. The ability to easily implement ODES on a laptop makes it accessible to a wide range of researchers, regardless of their computational resources or expertise. Its standardized, rule-based approach can reduce variability among researchers and speed up the classification process. ODES enables researchers to quickly and efficiently determine the estrous stage of female mice at multiple timepoints. This is particularly valuable in studies that require precise tracking of the estrous cycle. This is crucial in studies involving female mice and can lead to improved diagnostic precision in clinical settings for women. Despite its strengths, ODES faces challenges such as difficulty with unstained images and classification in clustered cell scenarios. These limitations can be mitigated with the expansion of the training dataset to incorporate unstained images and the shift to a newer or more complex model architecture.

## Acknowledgments

We would like to thank the Ross Lab, Verstegen Lab, and Correa Lab for providing vaginal cytology images. Funding sources for this work come from the Albert Einstein College of Medicine and the NIH DK118201 (RAR). In the preparation of this work, the authors used ChatGPT4 to debug the code used to train and run the model. The code is available on the Ross Lab Github link: https://github.com/rossrudolphlab/ODES_Object_Detection_For_Estrous_Staging

## Author Contributions

Conceptualization - BB, SG, RAR; Data curation - BB, SG, RAR; Formal analysis - BB; Funding - RAR; Methodology - BB, SG; Software - BB, SG; Validation - BB, SG; Visualization - BB, SG; Writing - BB, SG, RAR.

## Supplementary Figures

**Supplemental Figure 1:**
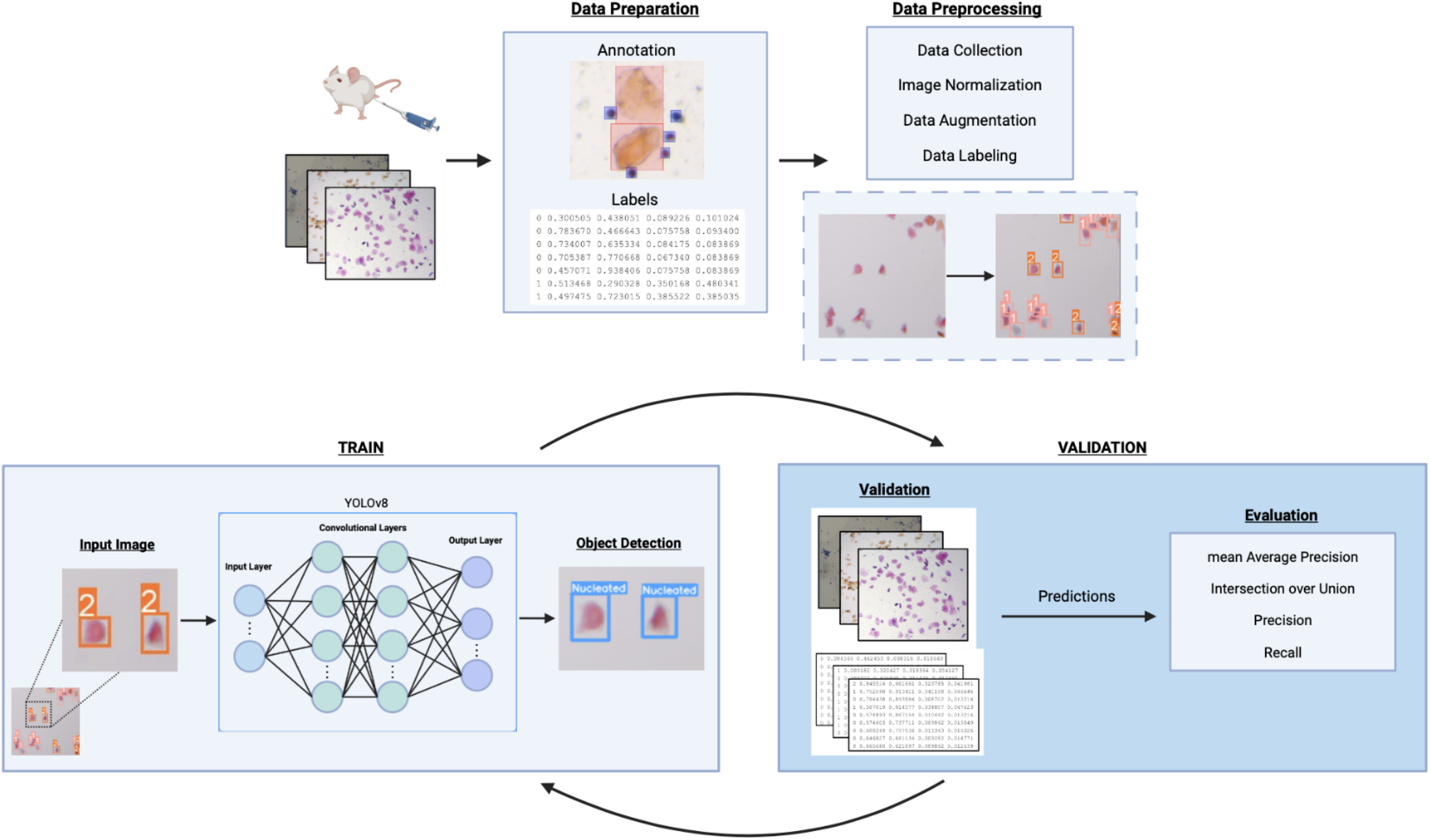
Overview of ODES Training. Schematic made with BioRender. Figure 10 illustrates the workflow diagram for supervised machine learning for the estrous cycle images of mice. A select portion of these images were annotated, marking individual cells within the image with their respective labels. The labels for each annotated object were then exported into a format that provides the coordinates for the bounding box (x center coordinate, y center coordinate, width, height). Afterward, the images were organized and placed into Yolov8 for training. The data was first preprocessed via data collection and normalized by image size and color by adjusting the pixel values to a consistent scale. The data was then augmented and labeled to complete the processing and begin the training. The input image is the starting point of the training, where it gets fed into the Yolov8 architecture. This architecture consists of complex machine learning techniques such as convolutional layers for feature extraction, optimization algorithms, and loss functions to adjust the model’s parameters. After it runs through the first batch of images and updates its weights, the model validates itself against a separate dataset, the validation dataset. The following metrics are used to evaluate its performance: Mean Average Precision (mAP), Intersection over Union (IoU), Precision, and Recall. IoU is used to measure the precision of the bounding box of the model by comparing the predicted bounding box to the correct bounding box [27]. Precision represents the model’s ability to avoid false positives, which is calculated by finding the ratio of true positives and total positive predictions. Recall calculates the ratio of true positives detected and all actual positives, measuring the model’s ability to detect all instances of a class. Average Precision (AP) is the area under the precision-recall curve that provides information on the model’s precision and recall performance. By extension, mAP calculates the average AP values across multiple object classes. This is useful in multi-class object detection scenarios to provide a comprehensive evaluation of the model’s performance. The training and validation sections of the model repeat in a loop for different batches of images until the set number of epochs or until there is no trend of improvement over a set number of epochs (patience).

**Supplemental Figure 2:**
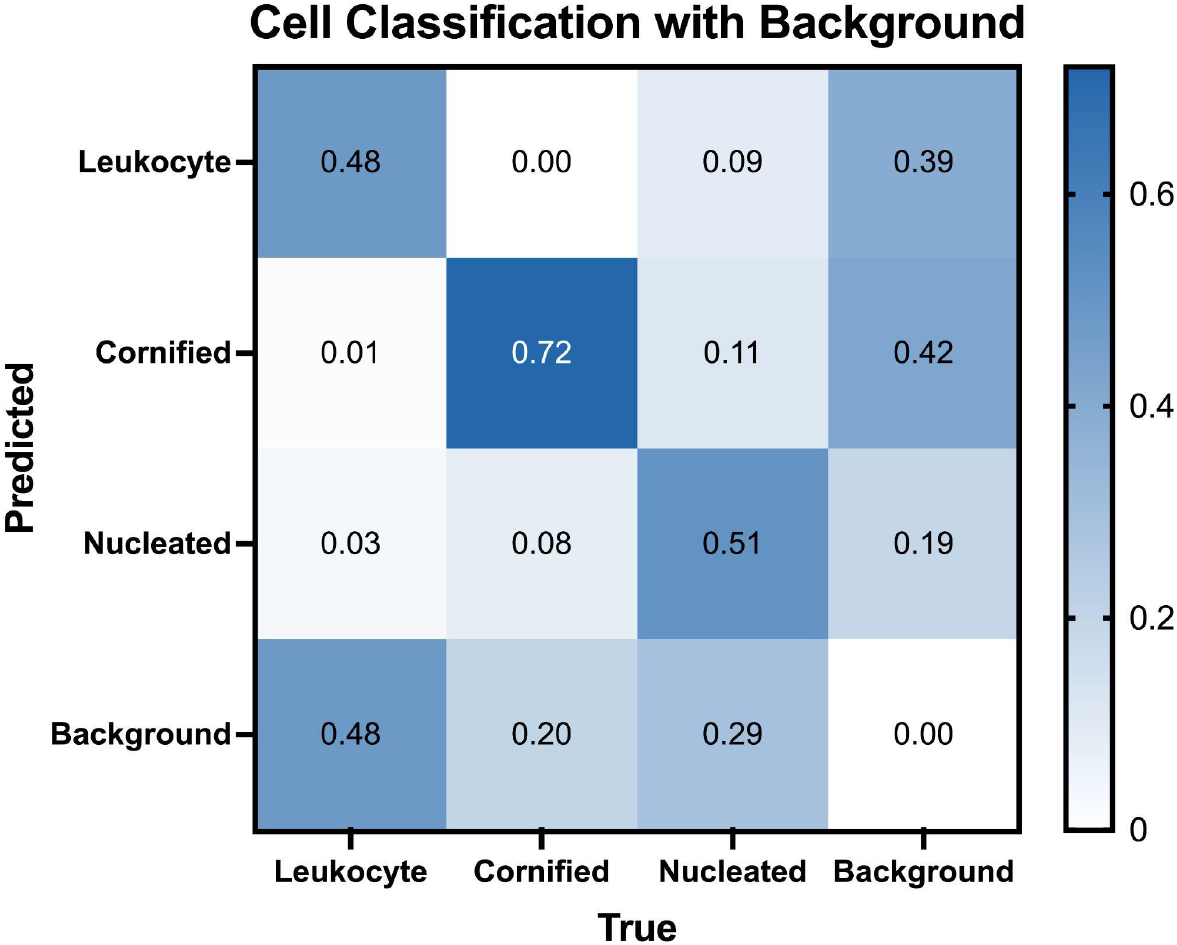
Comparative normalized confusion matrices for ODES cell classification with Background. With background classification, leukocytes were correctly identified 48% of the time but misclassified as background in 48% of instances. Cornified cells had a correct classification rate of 72%, with a 20% misclassification rate as the background. Nucleated cells were accurately classified at a rate of 51%, but 29% of instances were mistaken for background and an 8% rate of being mistaken for cornified cells.

**Supplemental Figure 3.**
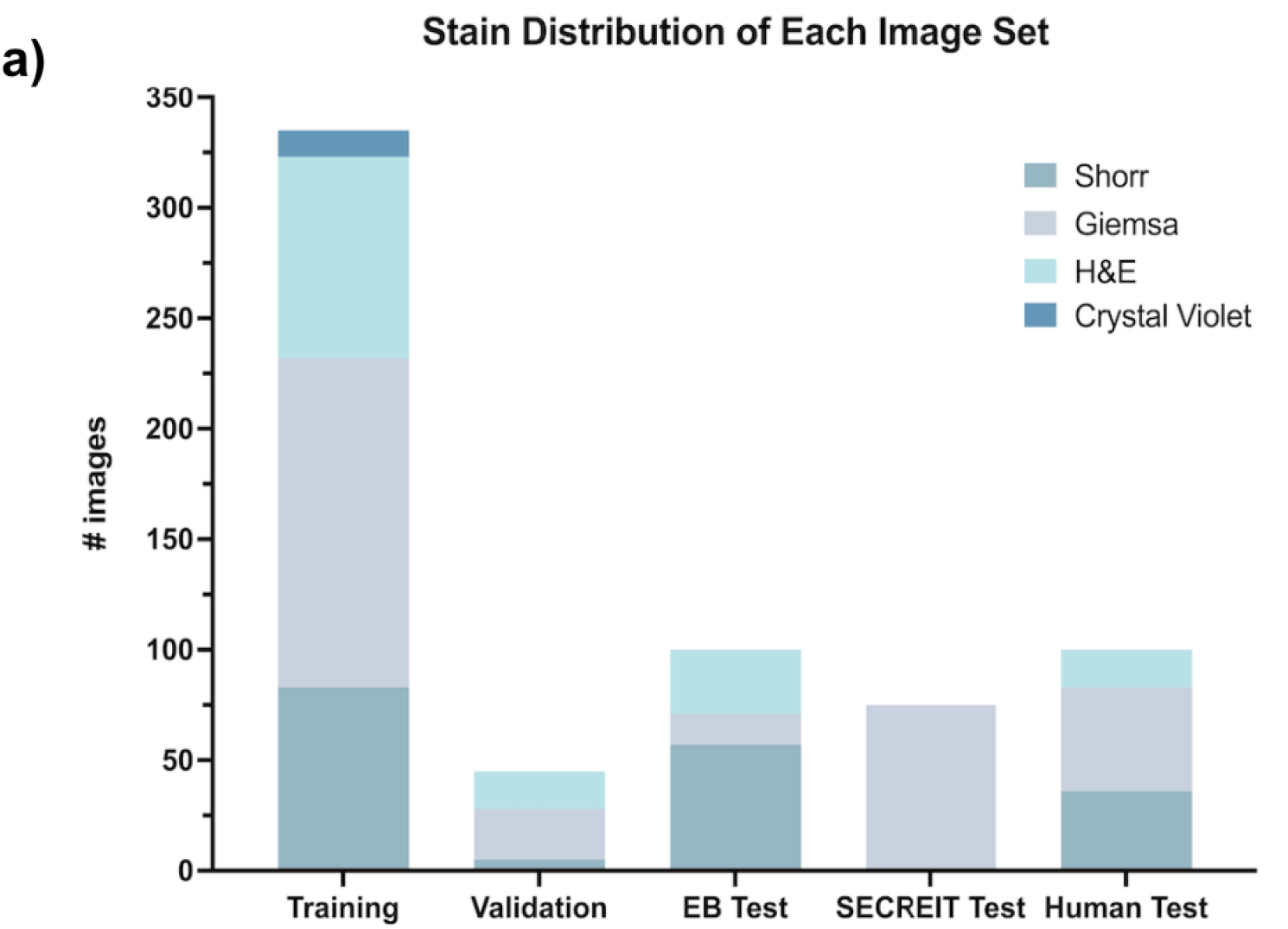
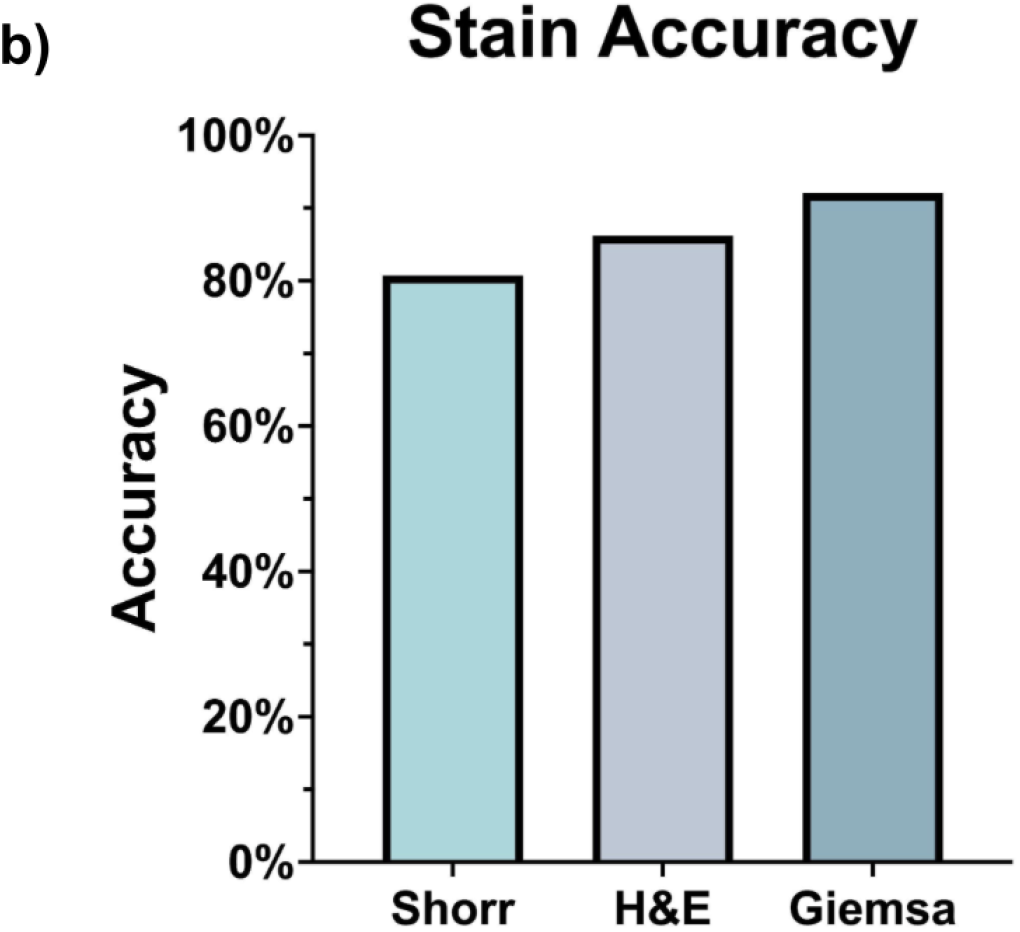
**a:** Distribution of vaginal cytology stains within the training and test sets. All images in the datasets were randomly placed. The training set included images with all four stains. The EstrousBank dataset predominantly contains images stained with the Shorr stain, supplemented by the Giemsa and H&E stains [11]. The SECREIT test was made exclusively of the Giemsa stained images from the SECREIT paper [10]. **b:** ODES accuracy for each stain in the test dataset. ODES performed best on images with a Giemsa stain however not to a degree of statistical significance. This test set included n=57 images with Shorr stain, n=29 images with H&E, and n=89 images with Giemsa divided across the four estrous cycle stages.

